# *μ*-rhythm extracted with personalized EEG filters correlates with corticospinal excitability in real-time phase-triggered EEG-TMS

**DOI:** 10.1101/426528

**Authors:** Natalie Schaworonkow, Pedro Caldana Gordon, Paolo Belardinelli, Ulf Ziemann, Til Ole Bergmann, Christoph Zrenner

## Abstract

Ongoing brain activity has been implicated in the modulation of cortical excitability. The combination of electroencephalography (EEG) and transcranial magnetic stimulation (TMS) in a real-time triggered setup is a novel method for testing hypotheses about the relationship between spontaneous neuronal oscillations, cortical excitability, and synaptic plasticity. For this method, a reliable real-time extraction of the neuronal signal of interest from scalp EEG with high signal-to-noise ratio (SNR) is of crucial importance. Here we compare individually tailored spatial filters as computed by spatial-spectral decomposition (SSD), which maximizes SNR in a frequency band of interest, against established local C3-centered Laplacian filters for the extraction of the sensorimotor *μ*-rhythm. Single-pulse TMS over the left primary motor cortex was synchronized with the surface positive or negative peak of the respective extracted signal, and motor evoked potentials (MEP) were recorded with electromyography (EMG) of a contralateral hand muscle. Both extraction methods led to a comparable degree of MEP amplitude modulation by phase of the sensorimotor *μ*-rhythm at the time of stimulation. This could be relevant for targeting other brain regions with no working benchmark such as the local C3-centered Laplacian filter, as sufficient SNR is an important prerequisite for reliable real-time single-trial detection of EEG features.

## 1. Introduction

Electroencephalography (EEG) provides access to neural dynamics on a millisecond timescale. In real-time EEG-triggered transcranial magnetic stimulation (TMS) it is possible to target specific brain states in applications such as personalized brain-stimulation [1]. However, as EEG is a mixture of different interacting sources, there is inherent ambiguity in inferring the brain state from the signal recorded with surface electrodes: the signal extracted depends not only on the source activity of interest but also on how the sensor channels are combined (i.e., the spatial filter) to maximally extract the source of interest while minimizing crosstalk from other sources and noise [2].

Whether the extracted signal corresponds to a functionally relevant brain state can be assessed by comparing TMS-evoked responses during different putative states. The relationship between TMS-evoked responses and ongoing oscillatory activity has previously been investigated with different methods regarding spatial filtering, e.g. in channel space [3–5], average over channel groups [6], with current source density [7], and with local spatial filters [1, 8]. These different approaches for defining brain states may explain some of the inconsistent results regarding the relationship between corticospinal excitability as measured by motor evoked potential (MEP) amplitude and features of EEG oscillations.

In this study, we computed a participant-specific spatial filter and a standard local filter (C3-centered Laplacian) to extract the sensorimotor *μ*-rhythm. Then we tested the dependence of corticospinal excitability on the phase of the extracted signal in a real-time triggered EEG-TMS setup.

The accuracy of our phase-estimation algorithm depends strongly on signal-to-noise ratio (SNR) [9]. Therefore, for computation of participant-specific spatial filters, we chose spatial-spectral decomposition (SSD) [10], a method designed to maximize the spectral power in a frequency band of interest while minimizing the power in neighboring (“noise”) frequency bands. SSD was also chosen because of its small set of parameters, the robust extraction of spatially localized oscillatory components with minor blurring [11], and its insensitivity to artefacts due to the usage of bandpass-filtered data, which enables fast computation during an experiment. The aim was to compare the degree of modulation of MEP amplitudes by the ongoing phase of the sensorimotor *μ*-rhythm (*μ*-phase) [1] as extracted by the two methods.

## 2. Materials and Methods

### 2.1 Participants

18 right-handed participants (4 male, 14 female, mean ± SD age: 24.99 ± 3.53 years, age range: 19–30), without a history of neurological disease or usage of CNS drugs, were selected from a pre-screened participant pool showing a clearly identifiable SNR in the *μ*-frequency band (8–13 Hz), with 5 dB above noise level. In this study, SNR is evaluated with a power spectrum from which the 1/f-component was subtracted [9]. Three participants were excluded after acquisition of resting EEG data because of excessive muscle artefacts, leaving 15 participants. Current TMS safety guidelines [12] were adhered to. All participants gave written informed consent prior to the experiment and tolerated the procedures without any adverse effects. The study protocol was approved by the ethics committee at the medical faculty of the University of Tübingen (protocol 716/2014BO2).

### 2.2 Experimental setup

#### 2.2.1 EEG and EMG recordings

The setup uses a combined EEG-TMS approach to trigger TMS pulses according to the instantaneous oscillatory phase of the extracted *μ*-rhythm. A 64-channel Ag/AgCl ring electrode EEG cap (EasyCap GmbH, Germany) was used, with an increased electrode density over the motor cortex (Figure 1A). A 24-bit amplifier was used for EEG and EMG recordings (NeurOne Tesla with Digital Out Option, Bittium Biosignals Ltd., Finland). EMG was recorded from relaxed right-hand muscles (right abductor pollicis brevis and first dorsal interosseous) using a bipolar belly-tendon montage.

**Figure 1:**
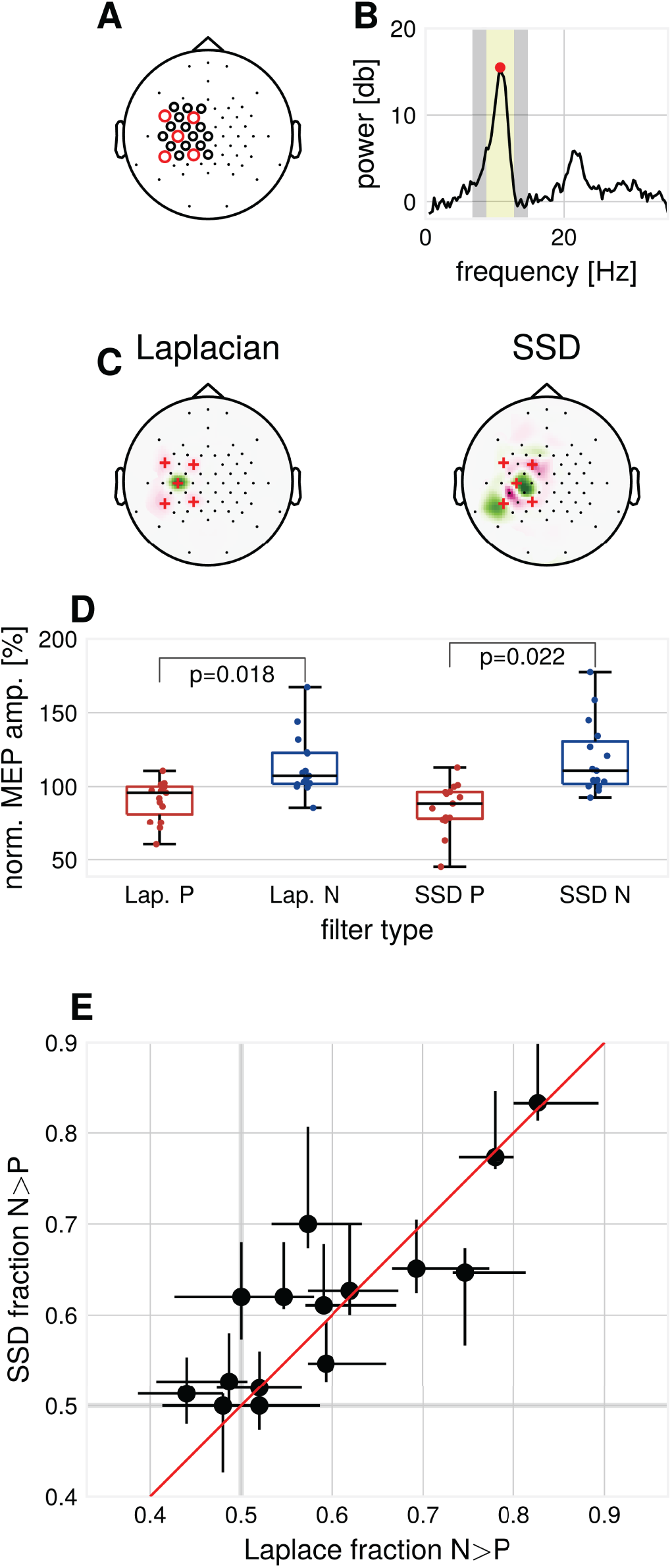
Methods and results: **(A)** EEG cap layout. Channels used for estimation of the individual spatial filters are marked with circles. Red circles indicate the channels used for determination of individual *μ*-peak-frequency. **(B)** Example 1/f-corrected spectrum used for determination of individual *μ*-peak frequency. Frequency bands used for the computation of SSD filters. Marked in yellow are the individual *μ*-peak frequency ±2 Hz, in grey the flanking noise frequency bands. **(C)** Example SSD spatial filter (right) computed from resting state EEG activity and the standard C3-centered Laplacian filter (left). **(D)** Group median MEP amplitudes for the respective filters, normalized by global median. p-values for Wilcoxon signed-rank test, multiple comparison corrected for the two types of filters, N=15. **(E)** Modulation of MEP amplitudes by *μ*-phase as assessed by the N/P-fraction for the respective filters, Laplace N/P-fraction and SSD N/P-fraction with 2.5 — 97.5^th^-percentile confidence intervals for each subject, N=15.

#### 2.2.2 TMS setup

Biphasic stimulation (AP-PA direction) was applied to the motor hotspot of the left primary motor cortex (coil position and orientation with maximal MEP amplitudes [13]) with a magnetic stimulator (Research 100, MAG & More GmbH, Germany) using a TMS double coil (PMD70-pCool, MAG & More GmbH, Germany). The target muscle was defined as the muscle with strongest responses at the lowest stimulator intensity. Resting motor threshold (RMT) was determined with a maximum likelihood PEST approach [14]. Neuronavigation (Localite GmbH, Germany) was used to maintain coil position.

#### 2.2.3 Real-time EEG-triggered brain stimulation

A real-time signal processing system was used to trigger TMS pulses according to the ongoing EEG (see [1]). The phase-detection algorithm, implemented in Simulink RealTime (Mathworks Ltd, USA, R2016a), was extended for processing signals filtered by two different spatial filters simultaneously. Single TMS pulses were triggered when the following conditions were met: (1) a minimum interstimulus interval (ISI) to the last pulse of 1.75 s was exceeded, (2) the instantaneous phase estimate for the signal filtered by the selected spatial filter fell within the specified target phase range, (3) a common oscillatory power threshold was exceeded simultaneously for the signals from both spatial filters in a 1024 ms sliding window. The power-threshold was monitored such that a median interstimulus interval of 2 s was attained. (4) The maximum peak-to-peak amplitude of the target muscle EMG during the last 500 ms signal was below a threshold of 50 *μ*V in order to prevent stimulation during muscle contraction.

#### 2.2.4 Computation of spatial filters

To minimize the influence of muscle artefacts and other non-stationarities, we used a reduced channel set to compute spatial filters (Figure 1A). Resting EEG data were band-pass filtered around the individual *μ*-peak frequency with ± 2 Hz, with 1 Hz width of the flanking frequency band defined as noise (Figure 1B). Two ipsilateral localized components were chosen from the resulting filter set (subsequently referred to as SSD#1 and SSD#2), according to the covariance-based spatial pattern [15]. SSD is invariant with respect to polarity. Therefore, the polarity of SSD spatial filters was aligned to minimize the mean phase difference to the Laplacian-filtered signal to maximize comparability. After computation, spatial filters were passed to the Simulink model for the phase-dependent TMS blocks (see Figure 1C for two example filters).

### 2.3. Experimental session

The experiment was structured as follows: (1) Eight minutes of eyes-open resting state EEG, used to compute participant-specific spatial filters for the main experiment. (2) Determination of RMT. (3) Three blocks of *μ*-phase-dependent stimulation with a fixed intensity of 112% RMT. In each block, four interleaved conditions were tested, triggering TMS pulses at surface *μ*-positive and *μ*-negative peaks for two spatial filters, respectively, referred to as P- and N-trials. The combination of the tested filters was fixed per block: 1) Laplacian vs. SSD#1, 2) Laplacian vs. SSD#2, 3) SSD#1 vs. SSD#2. 150 trials were acquired per condition resulting in a total of 1800 trials (2 phases × 2 filters × 150 trials × 3 blocks).

### 2.4. Data analysis and statistics

Data were analyzed with Matlab (Mathworks Ltd., USA, R2017b) and the BBCI toolbox [16]. EMG signals were high-pass filtered (Butterworth filter, order 4, cut-off 10 Hz). Peak-to-peak MEP amplitudes were determined within 20–60 ms after the TMS pulse. P- and N-trials were compared pairwise in the order as they appeared in the experiment (comparing the *i*^th^ N-trial with the *i*^th^ P-trial), calculating the N/P-fraction, the proportion of trials where MEP_N_ > MEP_P_. This procedure reduces the impact of slow time effects on absolute MEP size. The stronger the N/P-fraction deviates from 0.5, the stronger the observed phase-modulation. Confidence intervals were estimated with a bootstrap procedure by randomly shifting N- and P-MEP time courses against each other within a small window (shift drawn from uniform distribution [0, 25] trials, 10 000 iterations).

## 3. Results

### 3.1 Characteristics of individualized spatial filters

Contrary to pilot data on which the experimental protocol was based, generally, only one ipsilateral motor-component with high SNR could be extracted by SSD. Therefore, we focused on analyzing one specific session block for each participant. The block was selected according to similarity (measured by cosine distance) to the topography of the Laplacian spatial pattern.

While SSD improves SNR in contrast to using only sensor space data for channel C3 (mean SNR C3 channel = 12.08 ± 3.35 dB, mean SNR SSD = 15.27 ± 2.83 dB, Wilcoxon signed-rank test, *p* = 0.000122), no increase in SNR could be detected when comparing SNR for SSD and the Laplacian filter (mean SNR Laplacian = 14.14 ± 3.61 dB, mean SNR SSD = 15.27 ± 2.83 dB, Wilcoxon signed-rank test, *p* > 0.05). On resting state EEG data, we computed the mean phase shift between the Laplacian and the selected SSD filtered signal, bandpass filtered around the individual *μ*-frequency. On average over participants, a mean phase shift of 1.44° ± 3.93° was found, signifying a high correspondence in phase between Laplacian-filtered and SSD-filtered signals. We tested the phase accuracy of the real-time algorithm by passing the data through the Simulink model. A weak but significant decrease of the phase prediction error for SSD filters was detected (mean standard deviation of the phase prediction error: Laplacian 47.17°, SSD 44.25°, Wilcoxon signed-rank test, *α* = 0.05,*p* = 0.01).

### 3.2 Modulation of MEP amplitudes by *μ*-phase

Consistent with [1], for the Laplacian filter, we found larger MEP amplitudes for N-trials compared to P-trials (Wilcoxon signed-rank test, *α* = 0.05, *p* = 0.018, Figure 1D). Across participants, MEP amplitudes in N-trials were larger in 59.5% ± 11.8% (mean ± standard deviation) of trials compared to MEP amplitudes in successive P-trials. In terms of relative increase, median MEP amplitudes for N-trials were 1.33 ± 0.48 times larger compared to median MEP amplitudes for P-trials. For SSD filters, also a significant difference between median MEP amplitudes of P- and N-trials was found (Wilcoxon signed-rank test, *α* = 0.05, *p* = 0.022, Bonferroni-corrected for the two types of spatial filters).

We compared the N/P-fraction computed from MEP amplitudes for P- and N-trials for each type of spatial filter, see Figure 1E. The degree of observed modulation by *μ*-phase varied substantially between participants. On a group level, no significant difference between the N/P-fraction obtained by Laplacian filter stimulation and the individualized filter stimulation was detectable (Wilcoxon signed-rank test, *p* > 0.05). Also, at the individual participant level, no participant showed significant improvements with individualized filters when correcting for multiple comparisons (*χ*^2^-test of proportions, *p* > 0.05, Bonferroni-corrected for the number of tested particpants).

For one participant, two ipsilateral motor SSD-components could be extracted. Each filter led to a signal for which a significant difference between P- and N-trials could be detected. While one SSD filter showed no offset to the Laplacian filter, the other SSD filter produced a signal with an offset of 40.19°. For this participant, we also looked at the third block, in which both SSD-filters were directly contrasted against each other. No significant difference in the degree of modulation was found (*χ*^2^-test of proportions, *p* > 0.05).

## 4. Discussion

Brain-state dependent brain stimulation has the potential to increase the effectiveness and reliability of therapeutic protocols. Targeting a fixed brain state may reduce variability in the direction and degree of induced plasticity [1]. Extracting a functionally relevant brain state is therefore of interest. In this study, we computed individualized spatial filters using SSD for extraction of the sensorimotor *μ*-rhythm and compared this approach with a Laplacian filter. Using a real-time EEG-TMS setting, we tested whether and to what degree MEP amplitudes were modulated by the phase of the spatially filtered EEG signals. We found that it is feasible to compute individualized filters efficiently online during the experiment. However, using individualized filters yielded the same degree of observed phase-modulation as the benchmark Laplacian filter.

The accuracy of the real-time phase-detection algorithm strongly depends on SNR. Sufficient SNR is therefore a prerequisite for brain-state dependent stimulation. Filters which maximize SNR may be applied in situations where there is no clear pre-defined benchmark filter. In this study, the level of *μ*-SNR was relatively high. The potential SNR improvement obtained from using individualized filters may be larger for participants with lower SNR. This may enable the inclusion of a larger participant subset in studies where high SNR is an inclusion criterion, as higher SNR aids the phase-accuracy of the real-time algorithm.

The sole focus on SNR may, however, not be the optimal objective, as the phase of the extracted oscillation can vary strongly depending on scalp electrode position with respect to the cortical generators in a traveling wave manner [17, 18]. This is illustrated in Figure 2 with a single-subject example of oscillatory phase of the sensorimotor *μ*-rhythm. Depending on the selected central electrode of the local spatial filter, the phase is shifted between closely neighboring electrodes. For this study, participant-specific filters did usually not yield a systematic phase shift compared to the C3-centered Laplacian filter. Future studies should investigate methods for establishing correspondence of phase on the level of sensor space signals with the phase in the source space. This could yield insights about the cortical generators of local co-existing sensorimotor rhythms [19], as observed for one participant of our study. An open question yet is to what extent triggering on source level signals will increase the effective size of excitability modulation by phase for a wider participant range.

**Figure 2:**
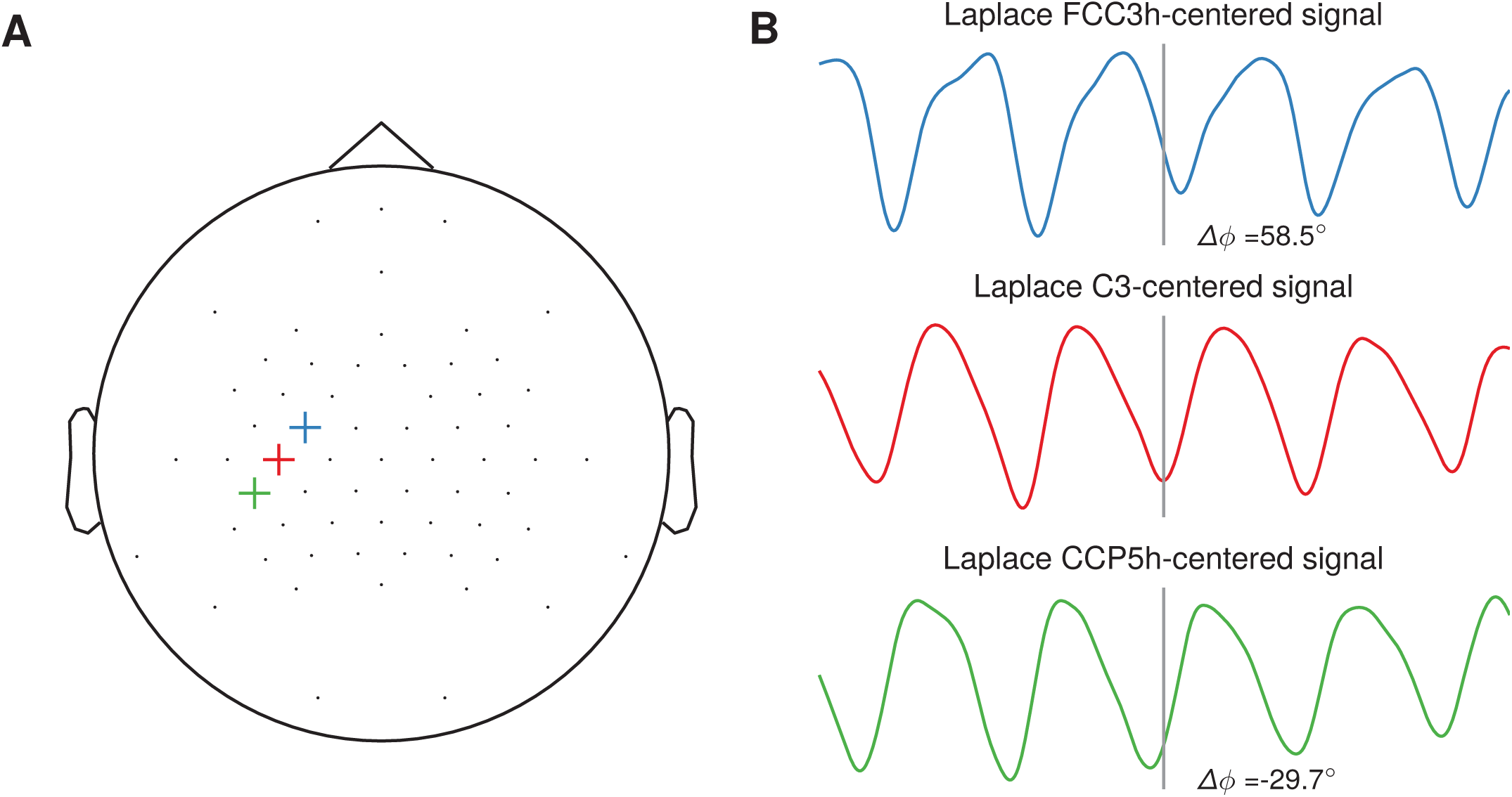
Illustration of phase shifts in sensor space. **(A)** Topography with three neighboring channels (FCC3h, C3, CCP5h) selected as center electrodes for the local spatial filter. **(B)** Resting-state EEG sensor space signals of one participant spatially filtered by a Laplacian filter centered on the selected electrode. The events are aligned to the troughs of the C3-centered Laplacian signal. A systematic phase shift is visible in the FCC3hand CCP5h-centered signal respective to the C3-centered Laplacian signal troughs, with 58:5° and –29:7°, respectively.

## Conflict of Interest Statement

The authors declare that the research was conducted in the absence of any commercial or financial relationships that could be construed as a potential conflict of interest.

## Author Contributions

All authors contributed to design and interpretation of experiments, manuscript revision, read and approved the submitted version. NS and PG performed data analysis. NS and CZ performed the experiments and wrote the first manuscript draft.

## Funding

NS, PG and CZ are supported by a EXIST Transfer of Research grant from the German Federal Ministry for Economic Affairs and Energy. CZ acknowledges support from Clinical Scientist Program at the Faculty of Medicine at the University of Tübingen. UZ acknowledges support from the German Research Foundation (grant ZI 542/7-1). TOB acknowledges support from the German Research Foundation (grant no. 362546008).

## Acknowledgments

We thank Anna Kempf and Dragana Galevska for help with participant coordination and experimental preparation. We thank Stasys Hiob for sharing MLS-PEST code.

## References

[1] C. Zrenner, D. Desideri, P. Belardinelli, U. Ziemann, Realtime EEG-defined excitability states determine efficacy of TMS-induced plasticity in human motor cortex, Brain Stimulation 11 (2018) 374–389.

[2] O. Hauk, M. Stenroos, A framework for the design of flexible cross-talk functions for spatial filtering of EEG/MEG data: DeFleCT, Human Brain Msapping 35 (2014) 1642–1653.

[3] L. Dugué, P. Marque, R. VanRullen, The phase of ongoing oscillations mediates the causal relation between brain excitation and visual perception., Journal of Neuroscience 31 (2011) 11889–11893.

[4] T. O. Bergmann, M. Mölle, M. A. Schmidt, C. Lindner, L. Marshall, J. Born, H. R. Siebner, EEG-guided transcranial magnetic stimulation reveals rapid shifts in motor cortical excitability during the human sleep slow oscillation, Journal of Neuroscience 32 (2012) 243–253.

[5] J. Keil, J. Timm, I. SanMiguel, H. Schulz, J. Obleser, M. Schönwiesner, Cortical brain states and corticospinal synchronization influence TMS-evoked motor potentials, Journal of Neurophysiology 111 (2014) 513–519.

[6] H. Mäki, R. J. Ilmoniemi, EEG oscillations and magnetically evoked motor potentials reflect motor system excitability in overlapping neuronal populations, Clinical neurophysiology 121 (2010) 492–501.

[7] B. Berger, T. Minarik, G. Liuzzi, F. C. Hummel, P. Sauseng, EEG oscillatory phase-dependent markers of corticospinal excitability in the resting brain, Biomedical Research International (2014).

[8] M. Thies, C. Zrenner, U. Ziemann, T. O. Bergmann, Sensorimotor mu-power is positively related to corticospinal excitability, Brain Stimulation (2018).

[9] N. Schaworonkow, J. Triesch, U. Ziemann, C. Zrenner, EEG-triggered TMS reveals stronger brain state-dependent modulation of motor evoked potentials at weaker stimulation intensities, Brain Stimulation (2018).

[10] V. V. Nikulin, G. Nolte, G. Curio, A novel method for reliable and fast extraction of neuronal EEG/MEG oscillations on the basis of spatio-spectral decomposition, NeuroImage 55 (2011) 1528–1535.

[11] S. Haufe, S. Dähne, V. V. Nikulin, Dimensionality reduction for the analysis of brain oscillations, NeuroImage 101 (2014) 583–597.

[12] S. Rossi, M. Hallett, P. M. Rossini, A. Pascual-Leone, S. of TMS Consensus Group, et al., Safety, ethical considerations, and application guidelines for the use of transcranial magnetic stimulation in clinical practice and research, Clinical neurophysiology 120 (2009) 2008–2039.

[13] P. Rossini, D. Burke, R. Chen, L. G. Cohen, Z. Daskalakis, R. Di Iorio, V. Di Lazzaro, F. Ferreri, P. Fitzgerald, M. George, M. Hallett, J. Lefaucheur, B. Langguth, H. Matsumoto, C. Miniussi, M. Nitsche, A. Pascual-Leone, W. Paulus, S. Rossi, J. C. Rothwell, H. R. Siebner, Y. Ugawa, V. F. Walsh, U. Ziemann, Non-invasive electrical and magnetic stimulation of the brain, spinal cord, roots and peripheral nerves: Basic principles and procedures for routine clinical and research application. An updated report from an I.F.C.N. Committee, Clinical Neurophysiology 126 (2015) 1071–1107.

[14] A. Mishory, C. Molnar, J. Koola, X. Li, F. A. Kozel, H. Myrick, Z. Stroud, Z. Nahas, M. S. George, The maximum-likelihood strategy for determining transcranial magnetic stimulation motor threshold, using parameter estimation by sequential testing is faster than conventional methods with similar precision, The Journal of ECT 20 (2004) 160–165.

[15] S. Haufe, F. Meinecke, K. Görgen, S. Dähne, J.-D. Haynes, B. Blankertz, F. Bießmann, On the interpretation of weight vectors of linear models in multivariate neuroimaging, NeuroImage 87 (2014) 96–110.

[16] B. Blankertz, L. Acqualagna, S. Dähne, S. Haufe, M. Schultze-Kraft, I. Sturm, M. Ušćumlic, M. A. Wenzel, G. Curio, K.-R. Müller, The Berlin brain-computer interface: progress beyond communication and control, Frontiers in Neuroscience 10 (2016).

[17] H. Suzuki, Phase Relationships of Alpha Rhythm in Man, Japanese Journal of Physiology 24 (1974) 569–586.

[18] P. L. Nunez, B. M. Wingeier, R. B. Silberstein, Spatial-temporal structures of human alpha rhythms: Theory, microcurrent sources, multiscale measurements, and global binding of local networks, Human Brain Mapping 13 (2001) 125–164.

[19] G. Pfurtscheller, C. Neuper, C. Andrew, G. Edlinger, Foot and hand area mu rhythms, International Journal of Psychophysiology 26 (1997) 121–135.

